# Spatial neglect after subcortical stroke: sometimes a cortico-cortical disconnection syndrome

**DOI:** 10.1101/2024.04.25.591066

**Authors:** Christoph Sperber, Hannah Rosenzopf, Max Wawrzyniak, Julian Klingbeil, Dorothee Saur, Hans-Otto Karnath

**Affiliations:** Center of Neurology, Division of Neuropsychology, Hertie-Institute for Clinical Brain Research, University of Tübingen, Tübingen; Department of Neurology, Neuroimaging Laboratory, University of Leipzig, Leipzig

**Keywords:** Thalamus, basal ganglia, white matter, hemineglect, attention, lesion network mapping

## Abstract

**Background and Objectives:** Spatial neglect is commonly attributed to lesions of a predominantly right-hemispheric cortical network. Although spatial neglect was also repeatedly observed after lesions to the basal ganglia and the thalamus, many anatomical network models omit these structures. We investigated if disruption of functional or structural connectivity can explain spatial neglect in subcortical stroke.

**Methods:** We retrospectively investigated data of first-ever, acute stroke patients with right-sided lesions of the basal ganglia (n = 27) or the thalamus (n = 16). Based on lesion location, we estimated i) functional connectivity via lesion-network mapping with normative resting state fMRI data, ii) structural white matter disconnection and iii) tract-wise disconnection of association fibres based on normative tractography data to investigate the association of spatial neglect and disconnection measures.

**Results:** Apart from very small clusters of functional disconnection observed in inferior/middle frontal regions in lesion-network symptom mapping for basal ganglia lesions, our analyses found no evidence of functional or structural subcortico-cortical disconnection. Instead, the multivariate consideration of lesion load to several association fibres predicted the occurrence of spatial neglect (p = 0.0048; AUC = 0.76), which were the superior longitudinal fasciculus, inferior occipitofrontal fasciculus, superior occipitofrontal fasciculus, and the uncinate fasciculus.

**Conclusion:** Disconnection of long (cortico-cortical) association fibres can explain spatial neglect in subcortical stroke. Like the competing theory of remote cortical hypoperfusion, this mechanism does not require the assumption of a genuine role for subcortical grey matter structures in spatial neglect.

## Introduction

Spatial attention deficits are a typical and debilitating consequence of right hemispheric stroke. The syndrome of spatial neglect, with an egocentric core deficit of an attentional deviation towards the ipsilesional side and neglect of contralesional objects ^1,2^, impacts stroke outcome and activities of daily living ^3,4^. A large research body was devoted to understanding the neural underpinnings of spatial neglect. The last two decades saw a shift from theories that focussed on single cortical loci in the right hemisphere, for example within the inferior parietal lobe ^5,6^, the superior or middle temporal lobes ^7,8^, and the inferior frontal lobe ^9,10^, to cortical network theories that unified these previous findings ^1,2,11^. This development was stimulated by advances in the accessibility of the human brain connectome and new insights into the impact of white matter disconnection on spatial neglect ^12–18^.

Infarcts in the thalamus and the basal ganglia were also repeatedly implicated in spatial neglect ^19–21^. A few theories incorporated such structures based on subcortico-cortical connections with cortical key regions of spatial neglect ^7,22–24^ or multimodal connectivity data ^25^, but subcortical grey matter structures only play a niche role in current network theories of spatial neglect. The functional role of subcortical grey matter structures is also challenged by an alternative explanation of spatial neglect after subcortical stroke that emphasizes the role of cortical hypoperfusion. In acute stroke with large vessel-occlusion, hypoperfusion typically goes beyond the core lesion area visible in diffusion-weighted MRI and includes areas where perfusion is sufficient to sustain tissue integrity but insufficient to sustain neural activity ^26,27^. In basal ganglia stroke, hypoperfusion and functional underactivation in cortical areas such as the superior temporal gyrus, the inferior parietal lobule, and the inferior frontal gyrus ^28^ can explain spatial neglect ^28,29^.

Yet another alternative account likewise dismisses the functional role of subcortical grey matter structures and explains spatial neglect after subcortical stroke with the disconnection of white matter fibres ^15,30,31^. Indeed, the disconnection of several major fibre bundles has been implicated in spatial neglect ^12,13,15,16,32^. These fibre bundles are located in the subcortical white matter directly adjacent to the thalamus and basal ganglia. Hence, even when a lesion of subcortical grey matter structures falls largely into the grey matter, it may still include damage to white matter fibre bundles which might suffice to cause spatial neglect through cortico-cortical disconnection.

In the current study, we aimed to re-evaluate the possible causes of spatial neglect in acute subcortical stroke using state-of-the-art analyses of indirect connectivity estimation. We studied patients whose lesions focussed on subcortical grey matter structures, without damage to cortical structures, and, at most, minor white matter damage. First, we used functional lesion-network symptom mapping, an approach to assess functional brain networks affected by focal structural lesions to gain insight into the possible role of subcortico-cortical connectivity. Second, we evaluated structural brain connectivity, including subcortico-cortical disconnection estimated based on normative connectome data. Third, we estimated the cortico-cortical disconnectome of each patient on the level of major fibre bundles to evaluate a possible role of cortico-cortical white matter disconnection.

## General methods

### Patient recruitment and behavioural assessment

We re-analysed data of stroke patients admitted to the Centre of Neurology at the University of Tübingen. Patients were retrospectively identified in datasets collected in previous studies ^8,20,33–35^. All patients had a first-ever unilateral cerebral stroke to the right hemisphere confirmed by CT or MRI imaging. Selection criteria were: 1) stroke affecting the basal ganglia or the thalamus, 2) no visible damage to cortical areas, and 3) no or only minor damage to white matter (average white matter lesion volume was 1.9 ± 2.25 cm³; range = 0.0–10.8 cm³). These criteria were first liberally checked by an automated search across available datasets by reference to the AICHA brain atlas ^36^. A second, visual evaluation of the neuroimaging of potentially suitable datasets resulted in a sample of 43 patients. Demographic and clinical data are shown in Table 1; lesion topographies in Figure 1. The study was approved by the local ethics board and has been performed in accordance with the revised Declaration of Helsinki. Patients or their relatives consented to the scientific use of the data.

**Table 1:** Clinical and demographic data of all patients.

| Patient Group | All | BG stroke, Neg+ | BG stroke, Neg- | Thalamus stroke, Neg+ | Thalamus stroke, Neg- |
| --- | --- | --- | --- | --- | --- |
| N | 43 | 10 | 17 | 4 | 12 |
| Age | 64.0 (12.7) | 66.5 (17.4) | 60.4 (11.1) | 72.0 (9.1) | 64.3 (11.1) |
| Sex (F/M) | 15/28 | 6/4 | 3/14 | 2/2 | 4/8 |
| Lesion to examination (days) | 4.7 (4.8) | 6.0 (6.1) | 4.4 (4.4) | 8.8 (6.2) | 2.6 (2.1) |
| Lesion to imaging (days) | 2.3 (4.1) | 3.3 (4.6) | 2.0 (2.4) | 9.5 (13.4) | 1.0 (1.8) |
| Neglect severity (CoC) | 0.16 (0.27) | 0.50 (0.31) | 0.01 (0.02) | 0.36 (0.22) | 0.00 (0.01) |
| Hemiparesis (%) | 77 | 88 | 64 | 100 | 78 |

Demographic data with ‘mean (standard deviation)’. BG = basal ganglia; Neg+ = patients with spatial neglect; Neg- = patients without spatial neglect; CoC = centre of cancellation (Rorden and Karnath, 2010).

**Figure 1:**
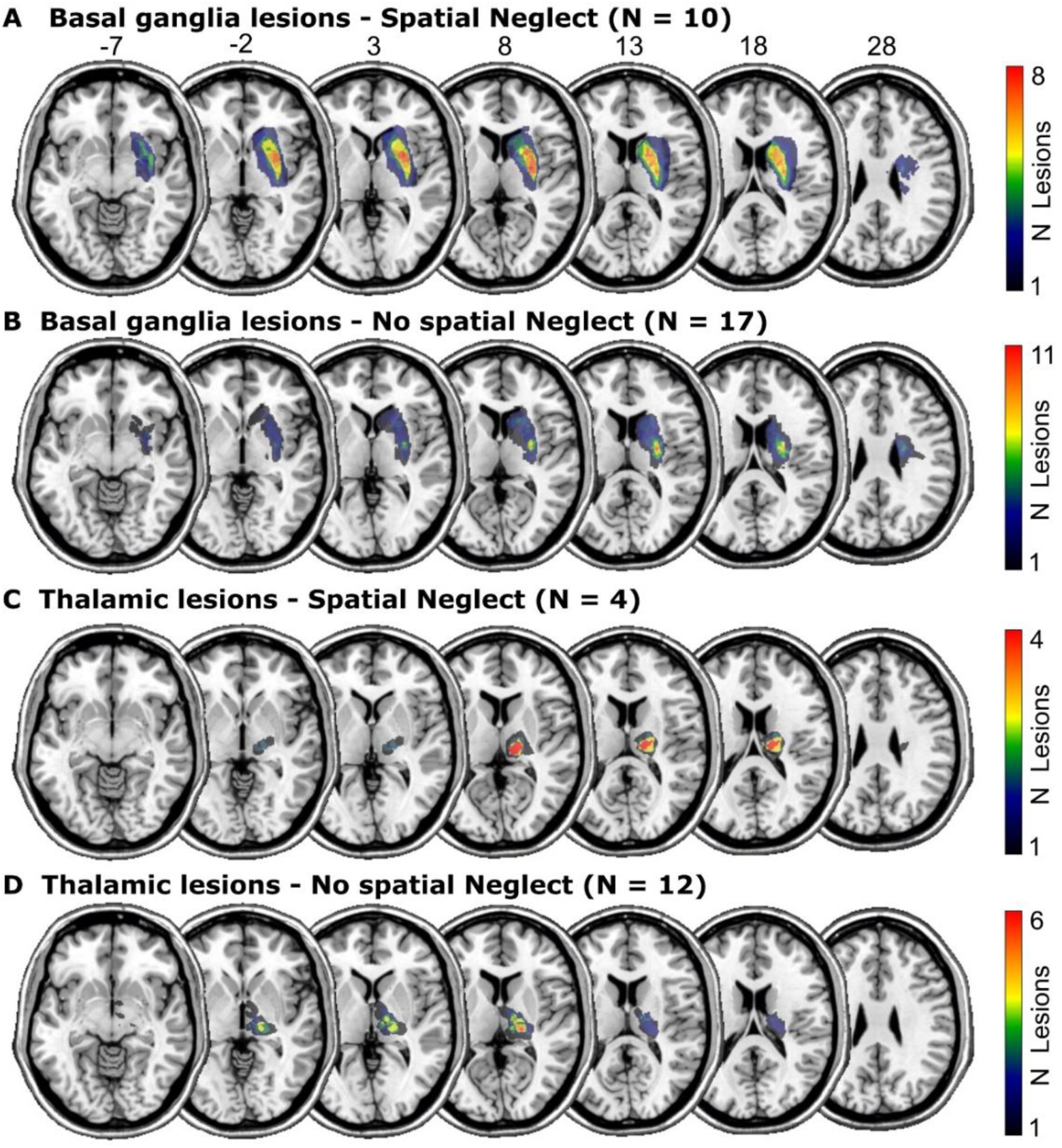
Lesion anatomy. Overlap topographies of structural lesion maps

Spatial neglect was assessed with pen and paper cancellation tests presented on 21 cm x 29.7 cm sheets of paper. In the letter cancellation task ^37^, the patients had to mark letters ‘A’ among distractor letters and, in the bells cancellation task ^38^, bell-shaped black icons among other similarly depicted objects. We evaluated the Centre of Cancellation (CoC; Rorden and Karnath, 2010), a continuous measure of spatial neglect in cancellation tests, which is predictive of typical neurological signs of spatial neglect. It ranges from −1 (maximum neglect of right stimuli) to 0 (symmetric test performance) up to 1 (maximum neglect of left stimuli). The average CoC score of both tests was used as the measure of spatial neglect severity. For analyses with binary data, patients were classified as suffering from spatial neglect if the CoC in at least one of the two cancellation tasks was above the empirically determined cut-offs ^39^. For 7 patients, only the measurement in one of the cancellation tests was available, which we then used as the sole measure.

### Imaging and structural lesion-behaviour mapping

Structural brain imaging was acquired by clinical CT (n = 30) or MRI (n = 13) on average 2.3 ± 4.1 days after stroke onset. In patients with an available MRI, diffusion-weighted imaging was used in the first 48 h after stroke onset and T2 fluid-attenuated inversion recovery imaging afterwards. Part of the data included in the present study reached back to times before digitalization of imaging data. For 25 of such older cases, lesions were manually drawn on axial slices of the ch2 MNI T1 template in MRIcron (see Karnath et al., 2004), and interpolated in the z-direction as described previously ^34^. For the remaining 18 cases, lesions were delineated semi-automatically on axial slices of the clinical scan using the Clusterize Toolbox ^40^. If possible, images were co-registered with high-resolution T1 MRI. Scans were warped to 1 x 1 x 1 mm³ MNI coordinates using age-specific templates in the Clinical Toolbox ^41^, situation-dependently either using cost-function masking or enantiomorphic normalisation. The normalisation parameters were then applied to the lesion masks.

### Lesion-network symptom mapping based on normative rs-fMRI data

We mapped the whole-brain functional connectivity of each lesion with ‘lesion-network mapping’ ^42^. In this approach, only the patients’ lesion masks are needed, and the brain-wide disturbance caused by a lesion is determined by reference to normative resting state functional MRI (rs-fMRI) of a healthy control group (Fig. 2). In short, the binary lesion mask of each patient is used as a seed in a rs-fMRI analysis. For each patient, the correlation of spontaneous resting brain activity within voxels inside the seed area is computed for all healthy control subjects for each voxel in the brain. This results in a single, full-brain lesion-network topography per patient, indicating both positive and negative functional connectivity with the lesioned area, which can be used within common statistical topographic approaches to map the brain-wide disruption of the functional connectome. The procedures followed previous studies ^43^, and utilised public rs-fMRI data of 100 healthy young subjects. The detailed procedures are described in the supplementary.

**Figure 2:**
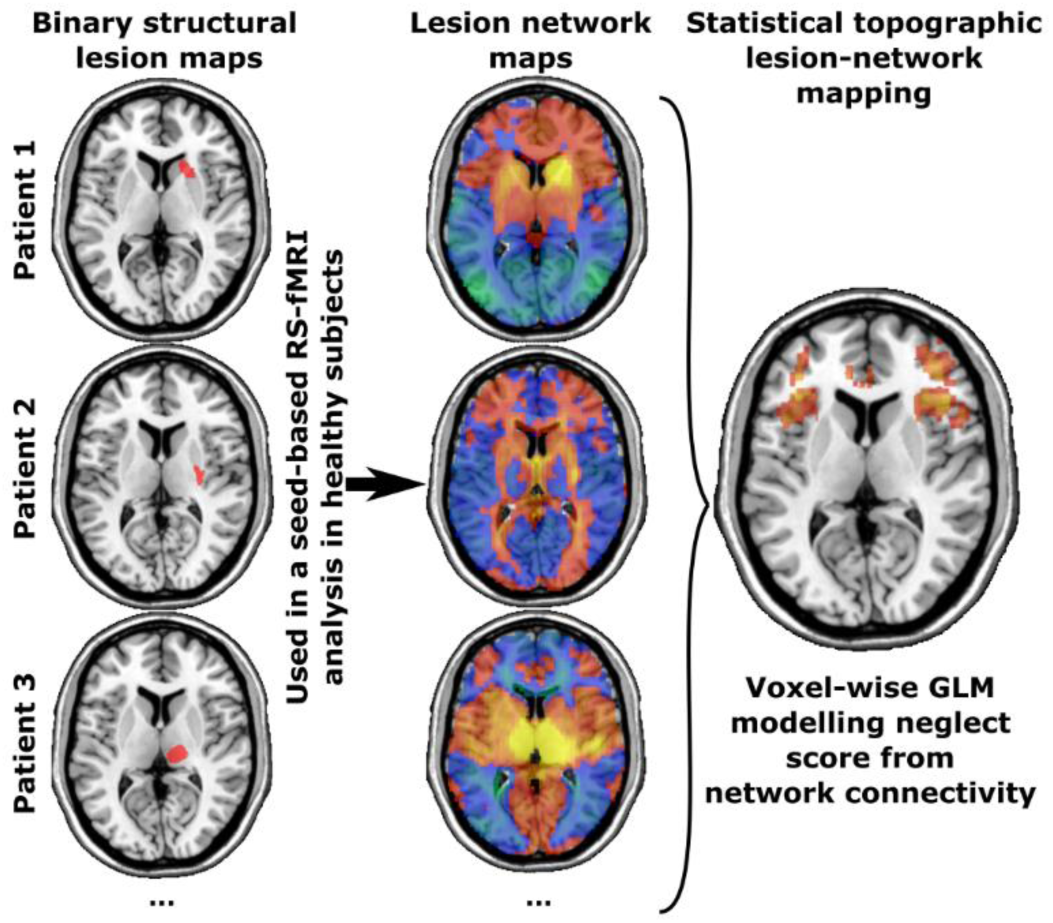
Concept lesion-network mapping. Design of the lesion network mapping analysis.

We utilised patient-wise maps of averaged Fisher-transformed Pearson correlations of rs-fMRI as lesion-network maps. For statistical analyses, we have masked the full-brain lesion-network maps to only include cerebral cortical grey matter areas with the AICHA brain atlas ^36^, henceforth termed cortical lesion-network maps. The masking step removed white matter areas and subcortical brain structures, i.e. the thalamus and basal ganglia, effectively removing areas where autocorrelation of resting state activity inside the lesion seed maps would inflate lesion-network connectivity.

### Estimated structural disconnectivity profiles

We indirectly estimated the lesion-induced disconnection of white matter fibres by reference to a healthy reference streamline connectome. Our method closely resembled the estimation of white matter disconnection in a popular toolkit ^44^, but extended this approach to the level of individual streamlines. We loaded all lesion masks as regions of interest to DSI studio (https://dsi-studio.labsolver.org/) and tracked streamlines based on the HCP842 ^45^ intersecting the lesion mask. The resulting set of streamlines included both streamlines that terminated in the lesion mask, i.e. disrupted connections of the subcortical structures themselves, as well as streamlines that passed the lesion mask, which included cortico-cortical streamlines.

### Tract-wise lesion load

We assessed the tract-wise lesion load of several major fibre tracts that were previously found to be implicated in spatial neglect ^12,16,18,33,46^. These included the superior longitudinal fasciculus (SLF), the inferior occipitofrontal fasciculus (IOF), the superior occipitofrontal fasciculus (SOF), and the uncinate fasciculus (UF). We defined the location of these fibre tracts with a histology-based probabilistic cytoarchitectonic fibre tract atlas ^47^. We created a binary region of interest for each tract by binarisation of the probabilistic map at p ≥ 0.3. The lesion load of a fibre tract was defined as the proportional overlap of the lesion map with this binarised map. While such overlap maps do not necessarily represent the the proportion of disconnected fibers, they were found (for motor deficits after damage to the corticospinal tract) to not only predict motor outcome but also that such maps performed better than lesion volume ^48^. In a control condition, we accounted for the full probabilistic information of the fibre tract atlas. We computed the weighted lesion load with the entire (non-thresholded) probabilistic fibre tract and the lesion map, whereas the lesion load of each voxel was weighted by its probability to be part of the fibre tract and summed up.

### Multivariate prediction analysis on tract-wise lesion load

We used a random forest classifier to evaluate if multivariate tract-wise lesion load can predict spatial neglect (methodologically similar to Mrah et al., 2022). We trained random forests with the scikit-learn package in Python to predict the binary spatial neglect diagnosis based on the four tract-wise lesion load measures. Each random forest contained 100 trees and, to prevent over-fitting, we limited the maximum depth to 3 and the minimum number of samples required to split an internal node to 5. We assessed the out-of-sample classifier performance in a looped cross-validation with 100 stratified splits of the total sample into 80 % training and 20 % test data. Classification performance was assessed in the test data by the area under the curve (AUC) of the receiver operator characteristic. We tested the classification performance against chance level by permutation testing. For 2500 times, we randomly shuffled the target variable and re-computed a random forest with the same procedures as for the original data. With this procedure, we assessed the distribution of the AUC under the null hypothesis and estimated the statistical significance of the original model. We also planned to repeat the multivariate random forest classification analysis with the indirectly estimated streamline-wise disconnectivity profiles aggregated for the major cortico-cortical fibre bundles based on the HCP842 ^45^. To do so, we derived patients’ tract disconnection measures using the lesion quantification toolkit ^44^.

## Experiment 1: Lesion network symptom mapping

### Statistical analysis

First, we used frequentist statistical parametric mapping to identify subcortico-cortical lesion-network connectivity related to spatial neglect after stroke to either the basal ganglia or the thalamus. We computed voxel-wise general linear models using NiiStat (https://github.com/neurolabusc/NiiStat) to test the association between spatial neglect severity as measured by average CoC scores and indirectly estimated connectivity in cortical lesion-network maps. Family-wise correction for multiple comparisons was implemented by maximum statistic permutation thresholding at p < 0.05 (two-tailed) with 10000 permutations. We interpreted the results with the rs-fMRI-based AICHA atlas ^36^.

Second, we replicated the analysis with a Bayesian approach with Bayes factor mapping to gain insights into evidence for the null hypothesis h_0_ and statistical power. We used the Bayesian Lesion-Deficit Inference toolbox ^50^ with a design equivalent to the frequentist analysis. Bayesian general linear models tested if an association between cortical lesion-network connectivity and spatial neglect exists in two-sided tests against the null hypothesis that no such association exists. For voxels without evidence for an association between spatial neglect and lesion networks, it can differentiate between the absence of evidence, i.e. a lack of statistical power, and evidence for the null hypothesis. We classified the degree of evidence provided by the mapped Bayes factors according to established conventions ^51^.

### Results

The frequentist analyses failed to find cortical lesion-network correlates of spatial neglect after correction for multiple comparisons both for basal ganglia and thalamic stroke.

For basal ganglia stroke, the Bayesian analysis (Figure 3A) was, as expected ^50^, more liberal and found scattered clusters of evidence for lesion network-neglect associations (i.e. Bayes Factors [BF] > 3, meaning that h1 is at least 3 times more likely than h0). However, the level of evidence was mostly only moderate and clusters were very small, creating largely non-interpretable results. We found a single small, potentially interpretable cluster in the right inferior and middle frontal gyrus (in regions 20 and 30 in the AICHA atlas), albeit maximum BFs did not exceed 12. On the other hand, due to the small sample size, the Bayesian analysis was unable to provide evidence in favour of the null hypothesis (BF < 1/3). Hence, we sub-differentiated BFs that suggested no evidence for any hypothesis (BFs > 1/3 and < 3) into anecdotal evidence for h1 (BF > 1 and < 3) and anecdotal evidence for h0 (BF < 1 and > 1/3; see Wagenmakers et al., 2018). Anecdotal evidence for h0 was found in the majority of tested voxels (74.1 %) and comprised large parts of the right cerebral cortex, including most of the temporal and parietal lobes.

**Figure 3:**
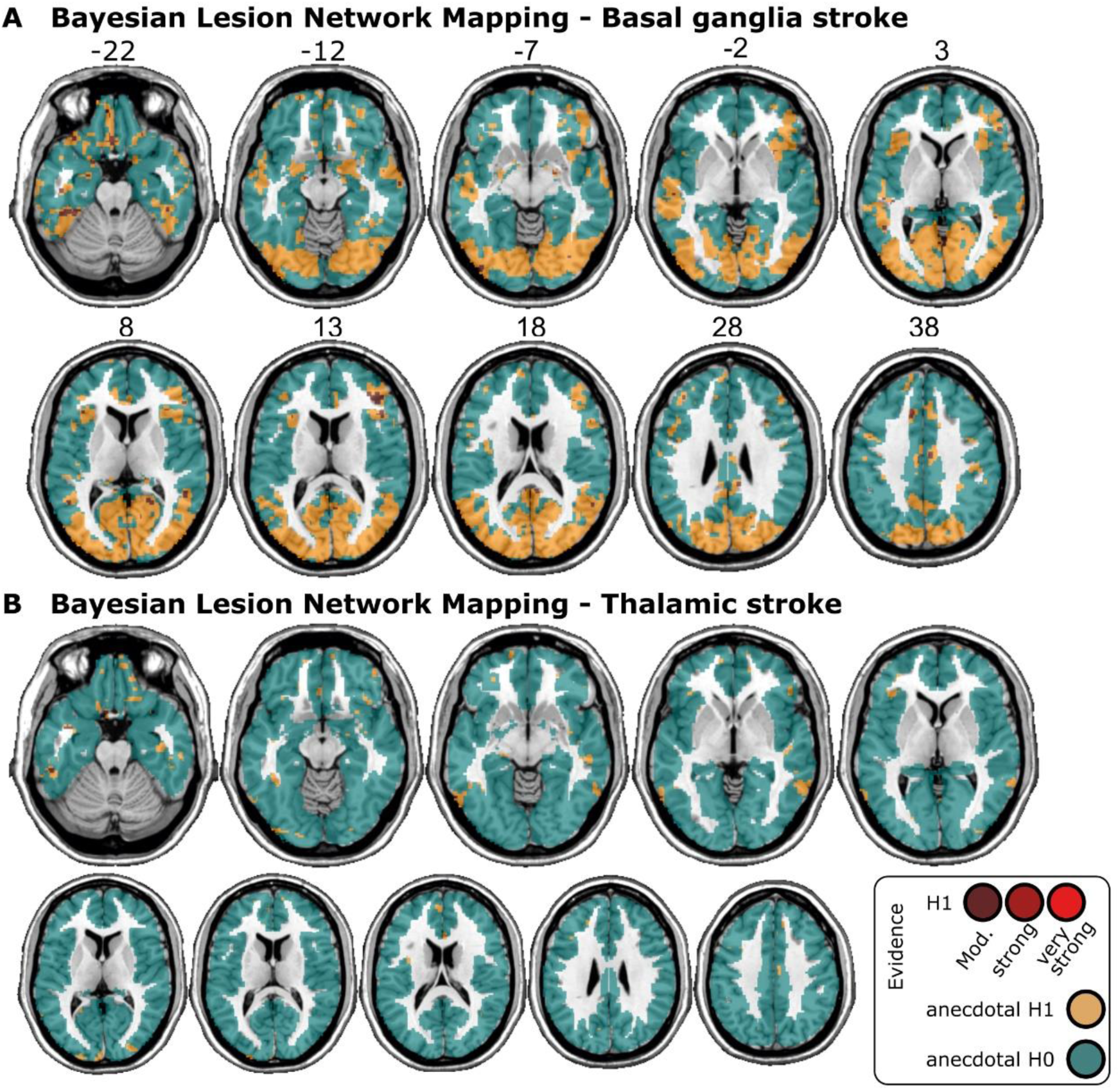
Results of the Bayesian lesion-network mapping analysis. **(A)** Statistical results of the Bayesian voxel-wise lesion-network analysis with normative rsfMRI data for basal ganglia stroke and **(B)** thalamic stroke. The analyses resulted in Bayes factors that weigh the evidence for the alternative hypothesis h_1_ (i.e. an association between the voxel-wise lesion-network connectivity and spatial neglect exists) and the null hypothesis h_0_ (i.e. no such association exists). For visualisation, we binned these Bayes factors according to common conventions (Wagenmakers et al., 2018) into evidence in favour of h_1_ (red; moderate BF > 3, strong BF > 10, very strong BF > 30), and inconclusive, anecdotal evidence for h1 (yellow; BF > 1) anecdotal evidence for h0 (turquoise; BF < 1). Due to the small sample sizes, the smallest Bayes factor was 0.35 (i.e. > 1/3), hence no moderate or stronger evidence for h0 was found.

For thalamic stroke, the Bayesian analysis found anecdotal evidence for h0 across the majority of tested voxels (97.9 %; Figure 3B) and no interpretable clusters of voxels with evidence for h1.

### Discussion

Voxel-wise univariate mapping failed to identify significant contributions of lesion network connectivity to spatial neglect after subcortical stroke. Moreover, Bayesian analyses partially provided anecdotal evidence for the absence of any association between lesion-network connectivity and spatial neglect in several known cortical key areas of spatial neglect. If anything, our analyses indicate functional connectivity of the basal ganglia with inferior/middle frontal due to very small clusters observed in these regions. However, due to the small sample sizes, we could not find at least moderate evidence in favour of the null hypothesis. In sum, lesion-network mapping did not find evidence for subcortico-cortical connectivity that could explain spatial neglect after stroke to the thalamus or basal ganglia.

## Experiment 2: Structural disconnectivity profile analysis

### Statistical analysis

We only tested streamlines disconnected in at least 5 patients, resulting in 88781 for basal ganglia lesions and 25873 streamlines for thalamic lesions. We tested the association of streamline disconnection with the mean CoC score with general linear models corrected by a permutation-based family-wise error correction with 10000 permutations at p < 0.05 (one-tailed). To assign significant streamlines to major fibre tracts we used the *recognize and cluster* function in DSI Studio.

### Results

In basal ganglia stroke, we found a small set of 12 streamlines in the corticostriatal tract (Figure 4A) where disconnection was significantly associated with spatial neglect. The identified streamlines appear to connect the putamen (AICHA region 366; Joliot et al., 2015) and a portion of the superior frontal gyrus (AICHA region 4 and 14) in the right hemisphere. In thalamic stroke (Figure 4B), disconnection of 141 streamlines in the right corticospinal tract, as well as a single streamline each from the right anterior thalamic radiation, right fornix, and the right dentato-rubro-thalamic tract were observed.

**Figure 4:**
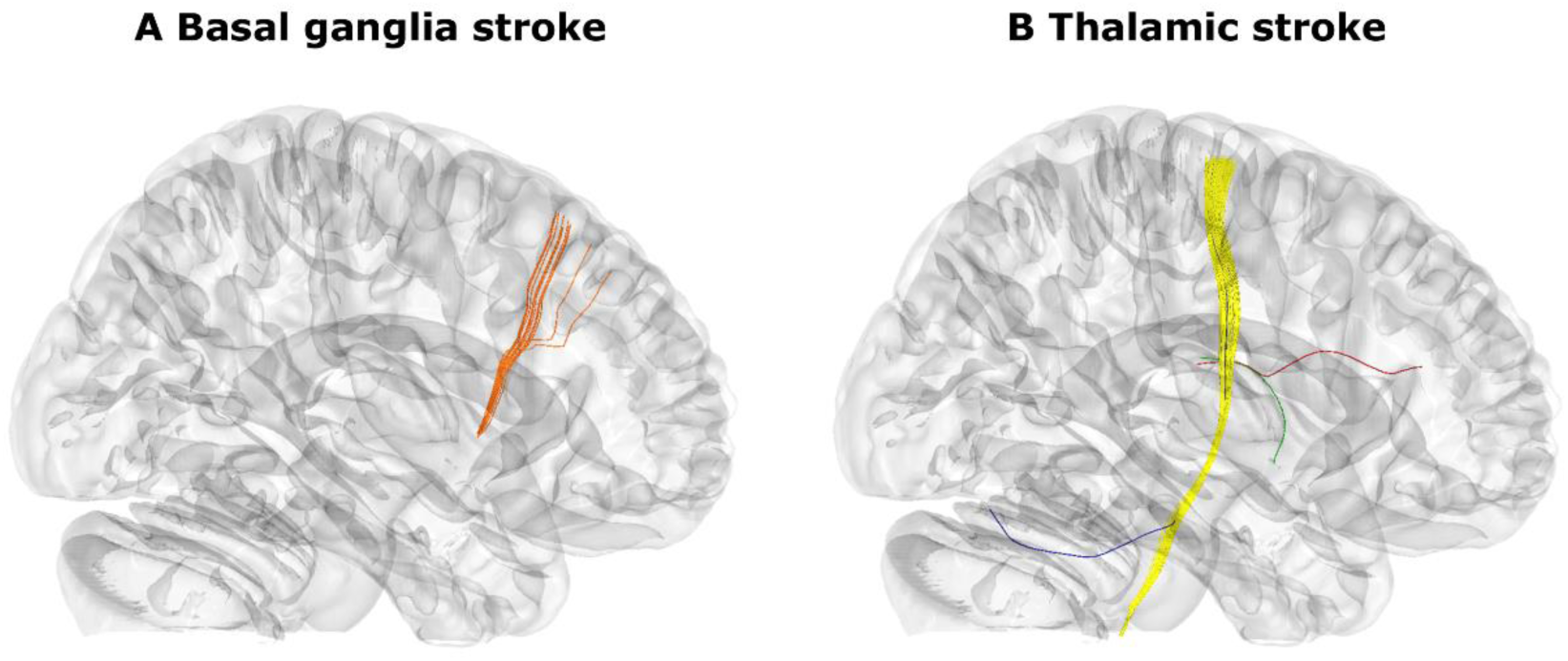
Streamline disconnections associated with neglect severity. **(A)** Significant disconnections of streamlines found in basal ganglia lesions. All fibres found were classified as part of the corticostriatal tract and are depicted in orange. **(B)** Significant disconnections in thalamic lesions. These included the corticospinal tract (yellow), anterior thalamic radiation (red), fornix (green) and dentatorubrothalamic tract (blue).

### Discussion

In basal ganglia stroke, cortical projections of fibres more often affected in neglect patients were located in the right superior frontal gyrus, i.e. more superiorly than those frontal regions commonly associated with spatial neglect (namely, the right ventrolateral frontal cortex [Karnath, 2009]). In thalamic stroke, the results were almost exclusively limited to the corticospinal tract. We consider this to be a coincidental finding unrelated to spatial neglect, but linked to the high rate of additional hemiparesis (Table 1) or to spatial inaccuracies that might have led to overlap with the external capsule. In summary, both analyses provided no convincing evidence of disconnectivity between subcortical and cortical regions to be causal for spatial neglect.

## Experiment 3: White matter tract-wise analysis

### Statistical analysis

With the third experiment, we investigated if spatial neglect can be explained by damage to white matter fibre tracts that pass in close proximity to the basal ganglia and thalamus. First, we assessed the association between tract-wise lesion load and spatial neglect. As the analysis focussed on cortico-cortical disconnection – independent of the subcortical lesion site in either the basal ganglia or thalamus – we performed analyses for all 43 patients combined. However, to improve comparability with analyses 1 and 2, we also report results for basal ganglia and thalamic stroke patients separately. Second, we used the random forest classifier to evaluate to what degree tract-wise lesion load can explain spatial neglect. In contrast to the simple tract-wise lesion load analysis, the random forest classification could not be repeated for the patient groups separately, since the overall sample size would be too small.

### Results

We found a lesion load in at least one of the four fibre tracts in 28 out of 43 patients. This was the case for 12 out of 15 patients with spatial neglect and 12 out of 29 patients without spatial neglect. This ratio was significantly larger in patients with spatial neglect (χ2(1, N = 43) = 3.88; p = 0.049). For all four fibre tracts, at least some patients with a lesion load were identified ranging from 13 to 23 patients (Figure 5). The rank correlation between the mean CoC score and the lesion load in each fibre tract ranged from τ = 0.20 to 0.27. However, while the correlation coefficients were significant for all fibres except the SLF, no results remained significant after Bonferroni correction. The correlation between the *total* lesion load across the four fibre tracts and the mean CoC was highly significant (τ = 0.31; p = 0.005; Figure 5), the correlation remained significant when only patients with damage to the basal ganglia were included (τ = 0.32; p = 0.023), but not when it was applied on the thalamic patient sample (τ = 0.24; p = 0.27) In a control analysis, we looked at the weighted lesion load that incorporated the full probabilistic information of the fibre tract atlas. This analysis closely replicated the correlation between lesion load and mean CoC (τ = 0.32; p = 0.003). The out-of-sample prediction accuracy of the random forest classifier was highly significant above chance (p = 0.0048; AUC = 0.76), meaning that the multivariate consideration of tract-wise disconnection can predict spatial neglect in subcortical stroke.

**Figure 5:**
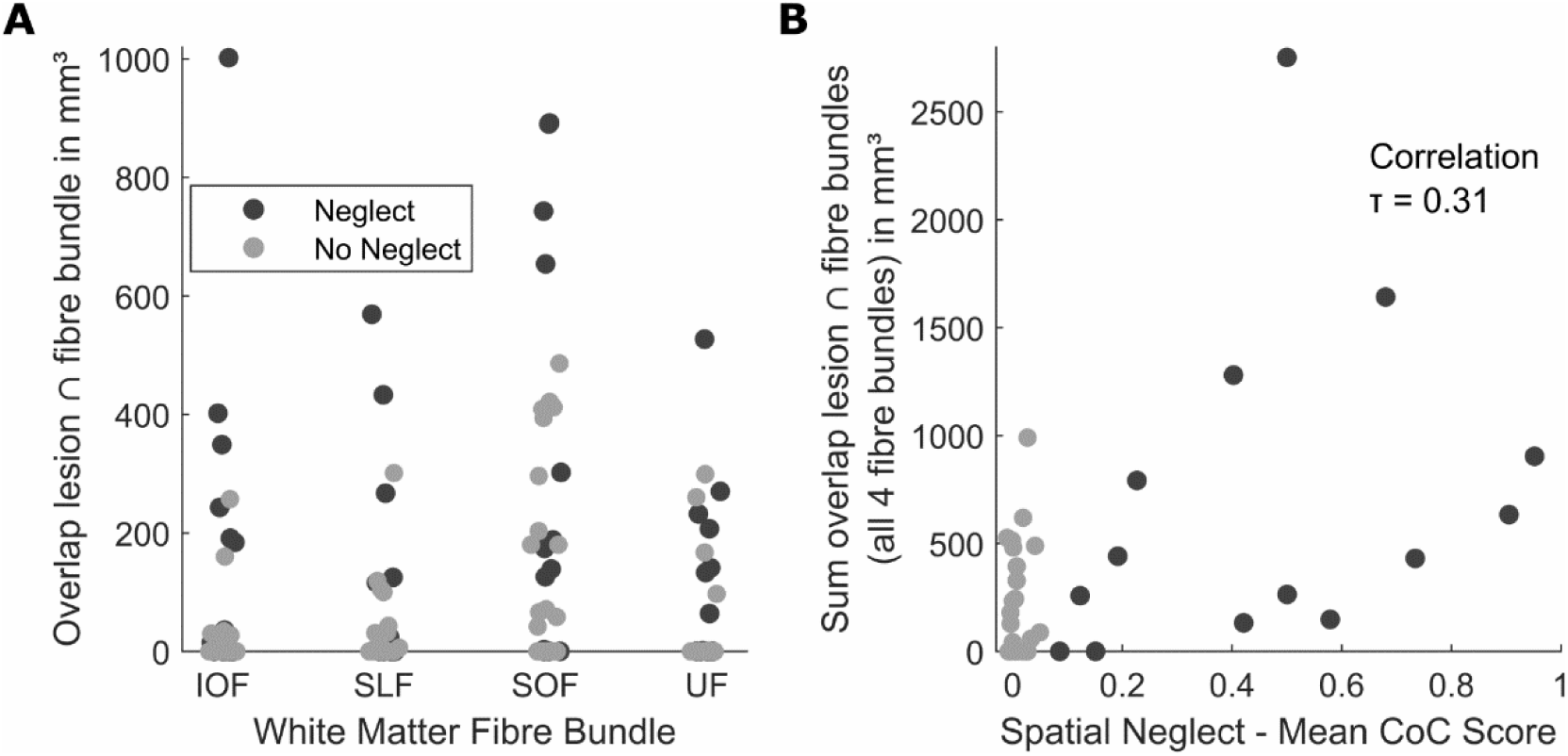
Results of the tract-wise analysis. **(A)** Scatter plots show the tract-wise lesion load for patients with and without spatial neglect, respectively **(B)** the relationship between total tract-wise lesion load and neglect severity.

We aimed to repeat the multivariate classification with the estimated streamline-wise disconnectivity across major cortico-cortical fibre bundles based on the HCP842 ^45^. However, not only the known neglect-related association fibre bundles (see above) were rarely or never disconnected, but association fibres in general. Disconnection of the SLF was found in only 2 patients. Only the IOF and the UF were more often found to be disconnected. Hence, we deemed the data to be unfit for multivariate model training.

### Discussion

The tract-wise lesion load analysis based on a probabilistic atlas of white matter bundles suggested disconnection of association fibres as a possible anatomical correlate of spatial neglect after subcortical stroke. Notably, it was the *multivariate* consideration of lesion load to several association fibres by the random forest classifier that predicted the occurrence of spatial neglect out-of-sample. This suggests that disconnection of different association fibres situated in proximity of the basal ganglia and thalamus could be a cause of spatial neglect in subcortical stroke.

## General Discussion

Our study investigated whether indirectly estimated functional or structural disconnection can explain spatial neglect in acute subcortical stroke to either the basal ganglia or the thalamus. Neither a functional analysis – i.e., lesion network mapping using normative resting-state fMRI – nor a structural analysis – i.e., the estimation of white matter disconnection using a normative streamline atlas – found convincing evidence for subcortico-cortical connectivity that could explain spatial neglect. However, we identified disconnection of cortico-cortical fibres that pass the subcortical structures in close proximity as a possible cause of spatial neglect.

The multivariate analysis of tract-wise lesion load based on a probabilistic atlas of white matter bundles suggested disconnection of cortico-cortical association fibres, namely the SLF, IOF, SOF, and the UF, as a possible neural correlate of spatial neglect after subcortical stroke. However, the same analysis of the disconnectivity profiles created with a non-probabilistic streamline atlas was not possible, as this method rarely or never captured disconnection of most of the potentially relevant fibres. A reason could be the marked variation in the spatial definition of white matter fibres across atlases, especially between atlases derived from histological versus from diffusion-tensor imaging data ^52^.

Since aphasia and spatial neglect in humans are caused by damage to somewhat homologue networks in opposing hemispheres, the state of research on aphasia following subcortical grey matter lesions might help to put our current findings into context. Aphasia after basal ganglia damage in the left hemisphere also remains controversial with heterogeneous findings and a lack of consensus on the involvement of specific structures ^53^. In line with the assumption of parallel mechanisms when considering our current results, white matter damage was found to be a potential explanation for the occurrence of aphasia in subcortical stroke ^54,55^. Other studies suggested, for example, left hemispheric thalamocortical functional disconnection as a possible mechanism ^56^. In our study, we did not find any results that would suggest similar mechanisms for spatial neglect after right-hemispheric thalamic stroke. However, we must note that our sample of right thalamic stroke cases (as opposed to right basal ganglia cases) was too small to draw or reject parallels with the explanation by Stockert and colleagues (2023).

Some anatomical network models have attributed a direct anatomical role to the basal ganglia, the thalamus, or both in the evocation of spatial neglect. These models were based on findings from lesion studies in subcortical stroke in comparison with existing knowledge about brain connections (e.g., Karnath et al., 2001; Parr and Friston, 2018). Other network models solely focused on cortico-cortical, intra- or interhemispheric networks and did not include any subcortical structures ^1,14^ In line with these theories, disconnection of long association fibres subserving a right fronto-temporo-parietal network have been found to underlie spatial neglect ^12,13,15,16,32^. The positive finding in our study was that cortico-cortical disconnection could explain spatial neglect, albeit this finding was only made with one out of two white matter-based analyses. The possibility of such a mechanism underlying spatial neglect in subcortical stroke adds − in addition to remotely induced hypoperfusion in intact cortical regions following lesions of subcortical grey matter structures ^28^ − yet another explanation that does not assume a direct role of subcortical structures in the spatial neglect network.

Given contradicting findings (e.g. Alves et al., 2022) and the possibility of multiple, parallel causal mechanisms, further studies are needed to understand the neural correlates of spatial neglect after subcortical stroke. This is of particular importance as null results in our study are of limited significance due to the relatively small sample size, as shown by the results of the Bayesian lesion network analysis. Bayesian statistics typically require larger samples to gather evidence for h0 than for h1^57^. Furthermore, even if a probable null effect existed, a possible role of subcortical structures in spatial neglect could not be excluded. A possible multitude of causes for spatial neglect in subcortical stroke – cortical hypoperfusion ^28,29^, cortico-cortical white matter disconnection, and a direct impact of damaged or disconnected subcortical structures – could statistically dilute an effect.

In summary, our study suggests that disconnection of cortico-cortical fibre bundles can explain spatial neglect after subcortical stroke. This explanation does not contradict the assumption that remote cortical hypoperfusion caused by subcortical damage causes spatial neglect ^28,29^. Both mechanisms seem plausible and, depending on the case, could underlie the occurrence of spatial neglect either separately or in combination. Apart from very small clusters of functional disconnection observed in inferior/middle frontal regions in lesion-network symptom mapping for basal ganglia lesions, our study found no evidence for the involvement of subcortico-cortical disconnection in the known network of spatial neglect. However, due to the small sample size in our study, we cannot exclude this possibility, and future studies with larger samples are needed to draw more precise conclusions on this question.

## Data availability

The data of the current study are not publicly available due to the data protection agreement approved by the local ethics committee and signed by the participants.

## CRediT author statement

**Christoph Sperber**: Conceptualization, Methodology, Data Curation, Formal Analysis, Writing - Original Draft;

**Hannah Rosenzopf**: Methodology, Formal Analysis, Writing - Original Draft;

**Max Wawrzyniak**: Methodology, Formal Analysis, Writing - Review and Editing;

**Julian Klingbeil**: Methodology, Formal Analysis, Writing - Review and Editing;

**Dorothee Saur**: Supervision, Writing - Review and Editing;

**Hans-Otto Karnath**: Conceptualization, Resources, Investigation, Data Curation, Supervision, Writing - Review and Editing.

## Abbreviations

CoC: Centre of Cancellation
IOF: inferior occipitofrontal fasciculus
rs-fMRI: Resting state functional magnetic resonance imaging
SLF: superior longitudinal fasciculus
SOF: superior occipitofrontal fasciculus
UF: uncinate fasciculus

## References

1. Corbetta M, Shulman GL. Spatial Neglect and Attention Networks. Vol 34.; 2011. doi:10.1146/annurev-neuro-061010-113731

2. Karnath HO, Rorden C. The anatomy of spatial neglect. Neuropsychologia. 2012;50(6):1010–1017. doi:10.1016/j.neuropsychologia.2011.06.027

3. Nijboer T, van de Port I, Schepers V, Post M, Visser-Meily A. Predicting functional outcome after stroke: The influence of neglect on basic activities in daily living. Front Hum Neurosci. 2013;7(MAY):1–6. doi:10.3389/fnhum.2013.00182

4. Moore MJ, Vancleef K, Riddoch MJ, Gillebert CR, Demeyere N. Recovery of Visuospatial Neglect Subtypes and Relationship to Functional Outcome Six Months After Stroke. Neurorehabil Neural Repair. 2021;35(9):823–835. doi:10.1177/15459683211032977

5. Mort DJ, Malhotra P, Mannan SK, et al. The anatomy of visual neglect. Brain. 2003;126(9):1986–1997. doi:10.1093/brain/awg200

6. Chechlacz M, Rotshtein P, Bickerton WL, Hansen PC, Deb S, Humphreys GW. Separating neural correlates of allocentric and egocentric neglect: Distinct cortical sites and common white matter disconnections. Cogn Neuropsychol. 2010;27(3):277–303. doi:10.1080/02643294.2010.519699

7. Karnath H-O, Ferber S, Himmelbach M. Spatial awareness is a function of the temporal not the posterior parietal lobe. Nature. 2001;411:950–953. doi:10.1016/S0010-9452(08)70654-3

8. Karnath HO, Berger MF, Küker W, Rorden C. The anatomy of spatial neglect based on voxelwise statistical analysis: A study of 140 patients. Cereb Cortex. 2004;14(10):1164–1172. doi:10.1093/cercor/bhh076

9. Husain M, Kennard C. Visual neglect associated with frontal lobe infarction. J Neurol. 1996;243(9):652–657. doi:10.1007/BF00878662

10. Committeri G, Pitzalis S, Galati G, et al. Neural bases of personal and extrapersonal neglect in humans. Brain. 2007;130(2):431–441. doi:10.1093/brain/awl265

11. Lunven M, Bartolomeo P. Attention and spatial cognition: Neural and anatomical substrates of visual neglect. Ann Phys Rehabil Med. 2017;60(3):124–129. doi:10.1016/j.rehab.2016.01.004

12. Thiebaut De Schotten M, Tomaiuolo F, Aiello M, et al. Damage to white matter pathways in subacute and chronic spatial neglect: A group study and 2 single-case studies with complete virtual “in vivo” tractography dissection. Cereb Cortex. 2014;24(3):691–706. doi:10.1093/cercor/bhs351

13. Thiebaut De Schotten M, Urbanski M, Duffau H, et al. Neuroscience: Direct evidence for a parietal-frontal pathway subserving spatial awareness in humans. Science (80- ). 2005;309(5744):2226–2228. doi:10.1126/science.1116251

14. Bartolomeo P, Thiebaut De Schotten M, Doricchi F. Left unilateral neglect as a disconnection syndrome. Cereb Cortex. 2007;17(11):2479–2490. doi:10.1093/cercor/bhl181

15. Doricchi F, Thiebaut de Schotten M, Tomaiuolo F, Bartolomeo P. White matter (dis)connections and gray matter (dys)functions in visual neglect: Gaining insights into the brain networks of spatial awareness. Cortex. 2008;44(8):983–995. doi:10.1016/j.cortex.2008.03.006

16. Karnath HO, Rorden C, Ticini LF. Damage to white matter fiber tracts in acute spatial neglect. Cereb Cortex. 2009;19(10):2331–2337. doi:10.1093/cercor/bhn250

17. Umarova RM, Reisert M, Beier TU, et al. Attention-network specific alterations of structural connectivity in the undamaged white matter in acute neglect. Hum Brain Mapp. 2014;35(9):4678–4692. doi:10.1002/hbm.22503

18. Toba MN, Migliaccio R, Batrancourt B, et al. Common brain networks for distinct deficits in visual neglect. A combined structural and tractography MRI approach. Neuropsychologia. 2018;115(October 2017):167–178. doi:10.1016/j.neuropsychologia.2017.10.018

19. Graff-radford NR, Damasio H, Yamada T, Eslinger PJ, Damasio AR. Nonhaemorrhagic thalamic infarction: Clinical, neuropsychological and electrophysiological findings in four anatomical groups defined by computerized tomography. Brain. 1985;108(2):485–516. doi:10.1093/brain/108.2.485

20. Karnath HO, Himmelbach M, Rorden C. The subcortical anatomy of human spatial neglect: Putamen, caudate nucleus and pulvinar. Brain. 2002;125(2):350–360. doi:10.1093/brain/awf032

21. Ten Brink AF, Biesbroek JM, Oort Q, Visser-Meily JMA, Nijboer TCW. Peripersonal and extrapersonal visuospatial neglect in different frames of reference: A brain lesion-symptom mapping study. Behav Brain Res. 2019;356:504–515. doi:10.1016/j.bbr.2018.06.010

22. Watson RT, Valenstein E, Heilman KM. Thalamic Neglect: Possible Role of the Medial Thalamus and Nucleus Reticularis in Behavior. Arch Neurol. 1981;38(8):501–506. doi:10.1001/archneur.1981.00510080063009

23. Parr T, Friston KJ. The Computational Anatomy of Visual Neglect. Cereb Cortex. 2018;28(2):777–790. doi:10.1093/cercor/bhx316

24. Saxena S, Keser Z, Rorden C, et al. Disruptions of the Human Connectome Associated With Hemispatial Neglect. Neurology. 2022;98(2):E107–E114. doi:10.1212/WNL.0000000000013050

25. Alves PN, Forkel SJ, Corbetta M, Thiebaut de Schotten M. The subcortical and neurochemical organization of the ventral and dorsal attention networks. Commun Biol. 2022;5(1). doi:10.1038/s42003-022-04281-0

26. Warach S, Dashe JF, Edelman RR. Clinical outcome in ischemic stroke predicted by early diffusion-weighted and perfusion magnetic resonance imaging: A preliminary analysis. J Cereb Blood Flow Metab. 1996;16(1):53–59. doi:10.1097/00004647-199601000-00006

27. Saur D, Buchert R, Knab R, Weiller C, Röther J. Iomazenil-single-photon emission computed tomography reveals selective neuronal loss in magnetic resonance-defined mismatch areas. Stroke. 2006;37(11):2713–2719. doi:10.1161/01.STR.0000244827.36393.8f

28. Karnath HO, Zopf R, Johannsen L, Berger MF, Nägele T, Klose U. Normalized perfusion MRI to identify common areas of dysfunction: Patients with basal ganglia neglect. Brain. 2005;128(10):2462–2469. doi:10.1093/brain/awh629

29. Hillis AE, Tuffiash E, Wityk RJ, Barker PB. Regions of neural dysfunction associated with impaired naming of actions and objects in acute stroke. Cogn Neuropsychol. 2002;19(6):523–534. doi:10.1080/02643290244000077

30. Doricchi F, Tomaiuolo F. The anatomy of neglect without hemianopia: A key role for parietal-frontal disconnection? Neuroreport. 2003;14(17):26–36. doi:10.1097/00001756-200312020-00021

31. Cha S, Jeong BC, Choi M, et al. White matter tracts involved in subcortical unilateral spatial neglect in subacute stroke. Front Neurol. 2022;13. doi:10.3389/fneur.2022.992107

32. Karnath H-O. A right perisylvian neural network for human spatial orienting. In: Gazzaniga MS, ed. The Cognitive Neurosciences IV. MIT Press; 2009:259–268.

33. Karnath HO, Rennig J, Johannsen L, Rorden C. The anatomy underlying acute versus chronic spatial neglect: A longitudinal study. Brain. 2011;134(3):903–912. doi:10.1093/brain/awq355

34. Sperber C, Karnath HO. Topography of acute stroke in a sample of 439 right brain damaged patients. NeuroImage Clin. 2016;10:124–128. doi:10.1016/j.nicl.2015.11.012

35. Wiesen D, Sperber C, Yourganov G, Rorden C, Karnath HO. Using machine learning-based lesion behavior mapping to identify anatomical networks of cognitive dysfunction: Spatial neglect and attention. Neuroimage. 2019;201(July):116000. doi:10.1016/j.neuroimage.2019.07.013

36. Joliot M, Jobard G, Naveau M, et al. AICHA: An atlas of intrinsic connectivity of homotopic areas. J Neurosci Methods. 2015;254:46–59. doi:10.1016/j.jneumeth.2015.07.013

37. Weintraub S, Mesulam M-M. Mental state assessment of young and elderly adults in behavioral neurology. In: Mesulam M-M, ed. Principles of Behavioral Neurology. F.A. Davis Company; 1985:71–123.

38. Gauthier L, Dehaut F, Joanette Y. The bells test: A quantitative and qualitative test for visual neglect. Int J Clin Neuropsychol. 1989;11:49–54. https://joanettelaboen.files.wordpress.com/2018/03/gauthier-l-dehaut-f-joanette-y-1989-version-finale-approuvecc81-yves.pdf

39. Rorden C, Karnath H-O. A simple measure of neglect severity. Neuropsychologia. 2010;48(9):2758–2763. doi:10.1016/j.neuropsychologia.2010.04.018.

40. De Haan B, Clas P, Juenger H, Wilke M, Karnath HO. Fast semi-automated lesion demarcation in stroke. NeuroImage Clin. 2015;9:69–74. doi:10.1016/j.nicl.2015.06.013

41. Rorden C, Bonilha L, Fridriksson J, Bender B, Karnath HO. Age-specific CT and MRI templates for spatial normalization. Neuroimage. 2012;61(4):957–965. doi:10.1016/j.neuroimage.2012.03.020

42. Boes AD, Prasad S, Liu H, et al. Network localization of neurological symptoms from focal brain lesions. Brain. 2015;138(10):3061–3075. doi:10.1093/brain/awv228

43. Wawrzyniak M, Klingbeil J, Zeller D, Saur D, Classen J. The neuronal network involved in self-attribution of an artificial hand: A lesion network-symptom-mapping study. Neuroimage. 2018;166:317–324. doi:10.1016/j.neuroimage.2017.11.011

44. Griffis JC, Metcalf N V., Corbetta M, Shulman GL. Lesion Quantification Toolkit: A MATLAB software tool for estimating grey matter damage and white matter disconnections in patients with focal brain lesions. NeuroImage Clin. 2021;30:102639. doi:10.1016/j.nicl.2021.102639

45. Yeh F-C, Panesar S, Fernandes D, et al. Population-Averaged Atlas of the Macroscale Human Structural Connectome and Its Network Topology. Neuroimage. 2018;178:57–68. doi:doi:10.1016/j.neuroimage.2018.05.027.

46. Vaessen MJ, Saj A, Lovblad KO, Gschwind M, Vuilleumier P. Structural white-matter connections mediating distinct behavioral components of spatial neglect in right brain-damaged patients. Cortex. 2016;77:54–68. doi:10.1016/j.cortex.2015.12.008

47. Bürgel U, Amunts K, Hoemke L, Mohlberg H, Gilsbach JM, Zilles K. White matter fiber tracts of the human brain: Three-dimensional mapping at microscopic resolution, topography and intersubject variability. Neuroimage. 2006;29(4):1092–1105. doi:10.1016/j.neuroimage.2005.08.040

48. Rickers E, Esser F, Rizor E, et al. Corticospinal tract lesion quantification: Distinct approaches and their association with motor impairment after stroke. Clin Neurophysiol. 2024;159:e24–e25. doi:10.1016/j.clinph.2023.12.064

49. Mrah S, Descoteaux M, Wager M, et al. Network-level prediction of set-shifting deterioration after lower-grade glioma resection. J Neurosurg. 2022;137(5):1329–1337. doi:10.3171/2022.1.JNS212257

50. Sperber C, Gallucci L, Smaczny S, Umarova R. Bayesian lesion-deficit inference with Bayes factor mapping: key advantages, limitations, and a toolbox. Neuroimage. 2023;271(March):120008. doi:10.1016/j.neuroimage.2023.120008

51. Wagenmakers EJ, Love J, Marsman M, et al. Bayesian inference for psychology. Part II: Example applications with JASP. Psychon Bull Rev. 2018;25(1):58–76. doi:10.3758/s13423-017-1323-7

52. de Haan B, Karnath HO. ‘Whose atlas I use, his song I sing?’ – The impact of anatomical atlases on fiber tract contributions to cognitive deficits after stroke. Neuroimage. 2017;163:301–309. doi:10.1016/j.neuroimage.2017.09.051

53. Radanovic M, Almeida VN. Subcortical Aphasia. Curr Neurol Neurosci Rep. 2021;21(12). doi:10.1007/s11910-021-01156-5

54. Alexander MP, Naeser MA, Palumbo CL. Correlations of subcortical CT lesion sites and aphasia profiles. Brain. 1987;110(4):961–988. doi:10.1093/brain/110.4.961

55. Damasio AR, Damasio H, Rizzo M, Varney N, Gersh F. Aphasia With Nonhemorrhagic Lesions in the Basal Ganglia and Internal Capsule. Arch Neurol. 1982;39(1):15–20. doi:10.1001/archneur.1982.00510130017003

56. Stockert A, Hormig-Rauber S, Wawrzyniak M, et al. Involvement of Thalamocortical Networks in Patients with Poststroke Thalamic Aphasia. Neurology. 2023;100(5):E485–E496. doi:10.1212/WNL.0000000000201488

57. van Ravenzwaaij D, Wagenmakers EJ. Advantages Masquerading as “Issues” in Bayesian Hypothesis Testing: A Commentary on Tendeiro and Kiers (2019). Psychol Methods. 2022;27(3):451–465. doi:10.1037/met0000415

