## supplementary for "Spatial neglect after subcortical stroke: sometimes a cortico-cortical disconnection syndrome"

### *Lesion-network mapping – detailed procedures*

Reference rs-fMRI data were taken from the human connectome project (van Essen et al., 2013) and included two resting-state sessions (with right-left and left-right phase encoding) of 15 minutes acquired with a gradient-echo echo planar imaging sequence (TR = 720 ms, 2 x 2 x 2 mm isotropic resolution). All images were already 'minimally preprocessed' when downloaded. This included gradient distortion correction, motion correction, distortion correction, normalization to MNI space, intensity normalization and bias field removal (Glasser et al., 2013). Functional images were smoothed (FWHM 5mm) and linearly corrected for motion parameters and their first derivate, and for mean white matter, cerebrospinal fluid and global signal (defined by tissue probability > 0.95). The remaining signal was band-pass filtered to retain frequencies between 0.01 and 0.08 Hz (Fox and Raichle, 2007). Motion Scrubbing was applied by calculating frame-wise displacement (Power et al., 2012) and discarding volumes with a value > 0.5mm. Two datasets were excluded because of heavy in-scanner motion with < 10 minutes of rs-fMRI after motion scrubbing in at least one session.

For the next level analyses, we used the binary normalized lesion map of each patient as a region of interest. Only grey matter portions of lesion maps were used in this step to ensure that included areas are subject to a meaningful BOLD signal. To this end, we utilised the grey matter tissue probability mask shipped with SPM12 at a probability threshold of > 0.1. Functional connectivity was computed as the Fisher-transformed Pearson correlation between the time series' first eigenvariate within the seed region and the time series of all remaining voxels. We calculated separate connectivity maps for the left-right and right-left phase encoding session. This procedure was repeated for each healthy subject and resulting topographies were averaged to produce a single topography ('lesion-network') per patient. We utilised patient-wise maps of averaged Fisher-transformed Pearson correlations as lesion-network maps. For statistical analyses, the full-brain lesion-network maps have been masked to only include cerebral cortical grey matter areas within the AICHA brain atlas (Joliot et al., 2015), termed cortical lesion-network maps.

***Supplementary References***

Fox, M.D., Raichle, M.E., 2007. Spontaneous fluctuations in brain activity observed with functional magnetic resonance imaging. Nat Rev Neurosci 8, 700–711. https://doi.org/nrn2201 [pii]\n10.1038/nrn2201

Glasser, M.F., Sotiropoulos, S.N., Wilson, J.A., Coalson, T.S., Fischl, B., Andersson, J.L., Xu, J., Jbabdi, S., Webster, M., Polimeni, J.R., Van Essen, D.C., Jenkinson, M., 2013. The minimal preprocessing pipelines for the Human Connectome Project. Neuroimage 80, 105–124. https://doi.org/10.1016/j.neuroimage.2013.04.127

Joliot, M., Jobard, G., Naveau, M., Delcroix, N., Petit, L., Zago, L., Crivello, F., Mellet, E., Mazoyer, B., Tzourio-Mazoyer, N., 2015. AICHA: An atlas of intrinsic connectivity of homotopic areas. J. Neurosci. Methods 254, 46–59. https://doi.org/10.1016/j.jneumeth.2015.07.013

Power, J.D., Barnes, K.A., Snyder, A.Z., Schlaggar, B.L., Petersen, S.E., 2012. Spurious but systematic correlations in functional connectivity MRI networks arise from subject motion. Neuroimage 59, 2142–2154. https://doi.org/10.1016/j.neuroimage.2011.10.018

Van Essen, D.C., Smith, S.M., Barch, D.M., Behrens, T.E.J., Yacoub, E., Ugurbil, K., 2013. The WU-Minn Human Connectome Project: An overview. Neuroimage 80, 62–79. https://doi.org/10.1016/j.neuroimage.2013.05.041
